# Size, shape and structure of insect wings

**DOI:** 10.1101/478768

**Authors:** Mary K. Salcedo, Jordan Hoffmann, Seth Donoughe, L. Mahadevan

## Abstract

The size, shape and structure of insect wings are intimately linked to their ability to fly. However, there are few systematic studies of the variability of the natural patterns in wing morphology across insects. We assemble a comprehensive dataset of insect wings and analyze their morphology using topological and geometric notions in terms of i) wing size and contour shape, ii) vein geometry and topology, and iii) shape and distribution of wing membrane domains. These morphospaces are a first-step in defining the diversity of wing patterns across insect orders and set the stage for investigating their functional consequences.

## INTRODUCTION

Of all the multicellular species on our planet, insects are the most speciose, with more than one million species^1^. Most are capable of flight owing to that remarkable evolutionary innovation, wings which themselves show a range of hierarchies of complexity, varying greatly in size and venation, stiffness and flexibility, pigmentation^6^ and flight behaviors^2–5^, while being subject to strong selective pressures by ecological niche specialization^7^.

From a physical perspective, insect wings are slender quasi two-dimensional membranes criss-crossed by a network of tubular veins. The patterns formed by veins can partition some wings into just a few domains and others into many thousands. The venation network allows for fluid and nutrient transport across the structure while providing a mechanical skeleton that stiffens the wing^3,8–10^. While phylogenetic analysis and geometric morphometrics have been deployed to understand the variation of and selection pressures on wing morphology^3,6,11–14^, their scope has been limited to a few species or orders at most. Here we complement these studies by using a set of simple quantitative measures to compare morphological variation in wing size, shape, and structure of insect wings across species, families and orders.

We start by assembling 555 wings drawn from 24 taxonomic orders, sampling representatives from nearly every extant insect order. We then deploy a range of geometrical and topological methods on these structures, and establish three complementary approaches to quantify wing size, shape, venation complexity and domain size and shape. Our dataset and analysis will, we hope, serve as a first step in functional and phylogenetic approaches to test hypotheses about wing evolution and physiology.

## MATERIALS AND METHODS

### Image collection and segmentation

Our dataset of wing images with representatives from most recognized insect orders is depicted in Fig. 1. We use a combination of original micrographs (hand-caught and donated specimens), a collection of scans from entomological literature (1840s - 1930s) sourced at the Ernst Mayr Library, Harvard University (Cambridge, MA) and online through the Biodiversity Heritage Library^15^(see SI for details).

**Figure 1.**
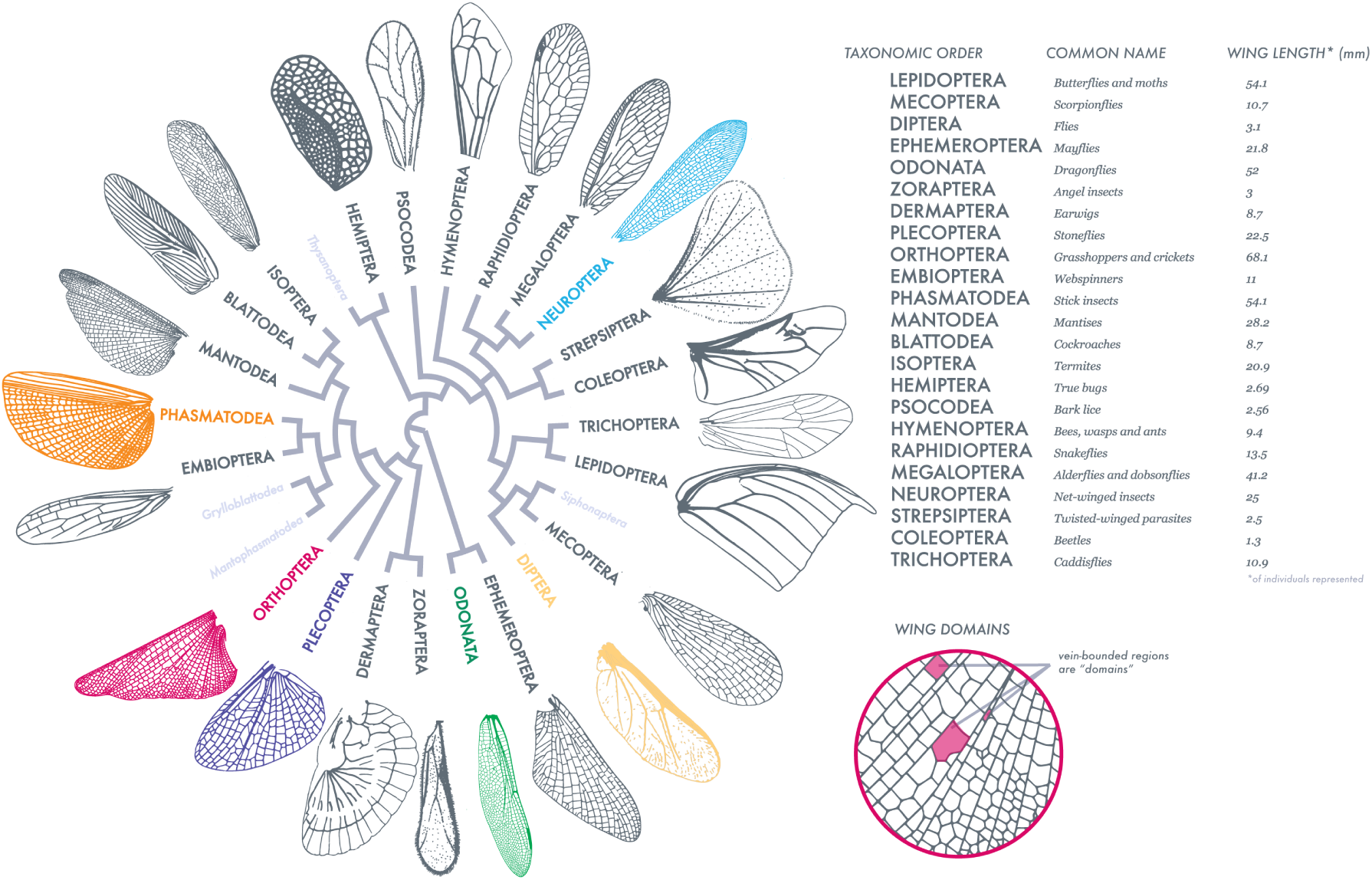
Wing taxonomy: size, shape and structure. Adapted from Misof et al.^26^, this order-level representation of insect wing size, venation patterns, shapes and wing domains (as defined in the figure on the right) exhibits the range and diversity of our sampling. Insect orders labeled in light gray are not sampled or characterized as wingless (full species list in SI). Scale bars are for wings represented on the phylogeny.

Entomological texts were chosen based on the quality and diversity of insects described and how well wing shape, venation, and morphological data were represented. This is not an unbiased sample of insects but we attempted to maximize diversity at the order level. By incorporating newly available data^16^, we obtained taxonomic coverage enriched for Odonata, a group with particularly complex wings. Insects are referred to by their common order-level names.

To quantitatively characterize an insect wing, we first segment a wing image using a Level-Set approach^16^ (see SI for details and dataset availability). Using this polygonal reconstruction, we are able to accurately and efficiently compute many geometric properties of an insect wing that is only possible with well-segmented data. For each wing, we also use the connectivity of neighboring vein domains and vertices to construct an adjacency matrix describing topological relationships between neighboring vertices.

## RESULTS

Since absolute wing size in insects is roughly correlated with body size^1^, we do not consider size directly. However, using these geometrical and topological datasets, we then calculate the basic geometric features of a wing: scaled venation length (Fig. 2a), scaled contour curvature (Fig. 2b), and scaled venation length (Fig. 2c). For each case, we normalize shape and wing contour to compare with a circle. We also studied the topology of venation using methods from network analysis (Fig. 2d). Finally, we studied the distributions of geometric domains, regions bound by veins, in terms of two simple statistical measures (Fig. 2e): 1) Circularity (*K*): the ratio of domain perimeter to the circumference of a circle with the same area and 2) Fractional area (*W*): the ratio of each individual domain area to the area of the entire wing. Together, these metrics serve as complementary features for quantifying the range of morphological characteristics of insect wings.

**Figure 2.**
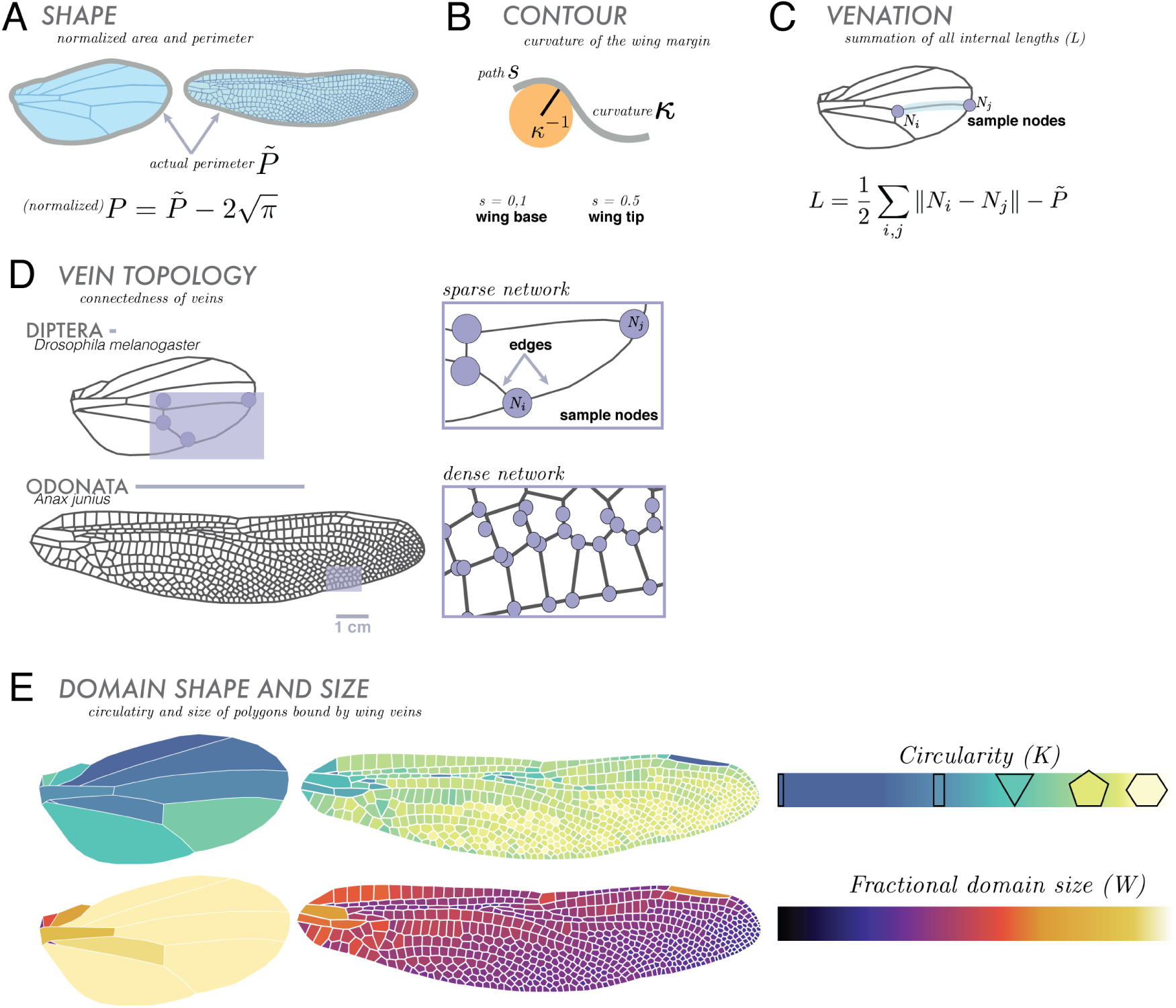
Wing morphometrics. We focus on broad comparative geometric and topological components, illustrated here using as examples Diptera (*Drosophila melanogaster*) and Odonata (*Anax junius*) wings. For geometric features, we analyze curvature, shape and area, and internal venation. **(A)** Contour, *κ*, is given by the radius of curvature or *κ*^*−*1^ (where *s* is arc length along the wing). **(B)** Shape: all wings are normalized to have an area equal to that of a circle with an area of unity (removing absolute size effects). Wing shape is characterized by its scaled perimeter, *P*, where 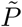 is the actual perimeter of the wing. **(C)** Venation is treated as a network, and quantified in terms of the sum of its total internal length, *L*, where *N_i_* and *N_j_* are representative nodes. We continue analyzing venation using topological measures where **(D)** the wing is a network of vein junctions (nodes) and the lengths of vein between them (edges). Lastly, we observe the geometries and distributions of vein domains. **(E)** Domains are characterized by their circularity (shape relative to that of a circle) and fractional domain size (domain area relative to area of entire wing).

### Wing shape, venation length and contour curvature

Our first morphometric feature characterizes all interior venation relative to wing contour, in terms of the scaled wing perimeter *P* (contour), and the scaled interior venation network, *L* (which excludes perimeter). Both of these geometric features are normalized by the square root of the wing area in order to isolate total vein length from overall wing size (Fig. 2a). Then, we define contour, the normalized perimeter,

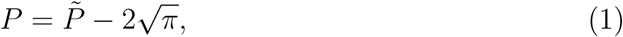

so that a circle with unit area would have *P* = 0. Additionally, *L*, the summation of all lengths of interior vein connectivity within the wing, yields the density of wing venation. In Fig. 3a, we plot the scaled wing contour length *L* against the venation contour *P*, noting that a wing at the origin (0, 0) corresponds to a circular wing without any internal venation. We see that wings with dense venation (e.g. locust, Orthoptera) occupy the upper middle/right sections of the graph and wings with sparse venation (e.g. fruit fly (Diptera)) occupy the lower/left regions of the morphospace.

**Figure 3.**
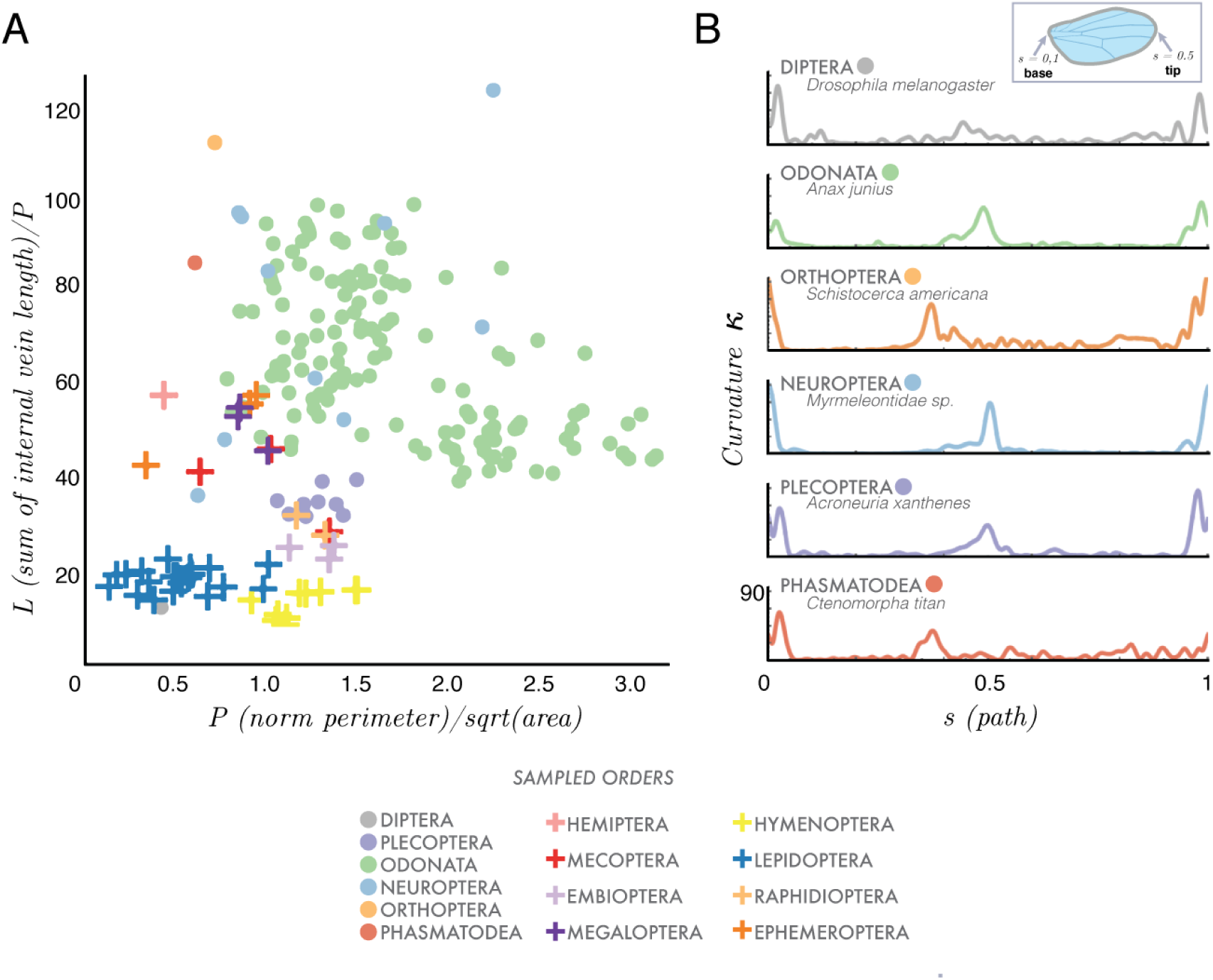
Wing shape, wing contour and internal venation. Comparison of three geometric traits (of all sampled orders) where **(A)** contour is defined using a scaled curvature (*κ*) (scaled by total perimeter *P*) as a function of arc length, *s*, where the wing base is *s* = 0 and the wing tip is *s* = 0.5. **(B)**. The total sum of all internal vein lengths *L* (scaled by the perimeter *P*), as a function of the normalized perimeter *P* (scaled by the square root of the area of the wing, see text) characterizes venation density. Species (per insect order) are represented by either circles or crosses.

Our second morphospace characterizes wing shape complexity treated in terms of its boundary curvature *κ*(*s*) as a function of the arc-length distance from the wing hinge. In Fig. 3b, we show a plot of wing curvature varying from wing base (*s* = 0 and *s* = 1) to wing tip (*s* = 0.5) for representative wings from six orders (top to bottom): *Diptera, Plecoptera, Odonata, Neuroptera, Orthoptera, Phasmatodea*. There seems to be no particular pattern to the distribution of curvature that we can discern.

### Wing vein topology

Wing venation forms a physical network, with the intersection of veins as nodes (Fig. 4). We use tools from network analysis^17,18^ to clustering the network into communities quantifying a third major trait of a wing: a topological measure of the complexity of venation patterns. We start by building an unweighted symmetric adjacency matrix, *A*, where every node_*ij*_ = node_*ji*_ (see Fig. 2d). To partition a wing network into clusters or communities^19^, we use the maximum modularity measure, which compares a given network to a randomly generated network and is maximized by a partition factor^20^ (see SI for other network versions comparison with more methods). This allows us to determine the number and size of clusters, each of which indicates a higher density of internal connections within a group of nodes, relative to connections across clusters.

**Figure 4.**
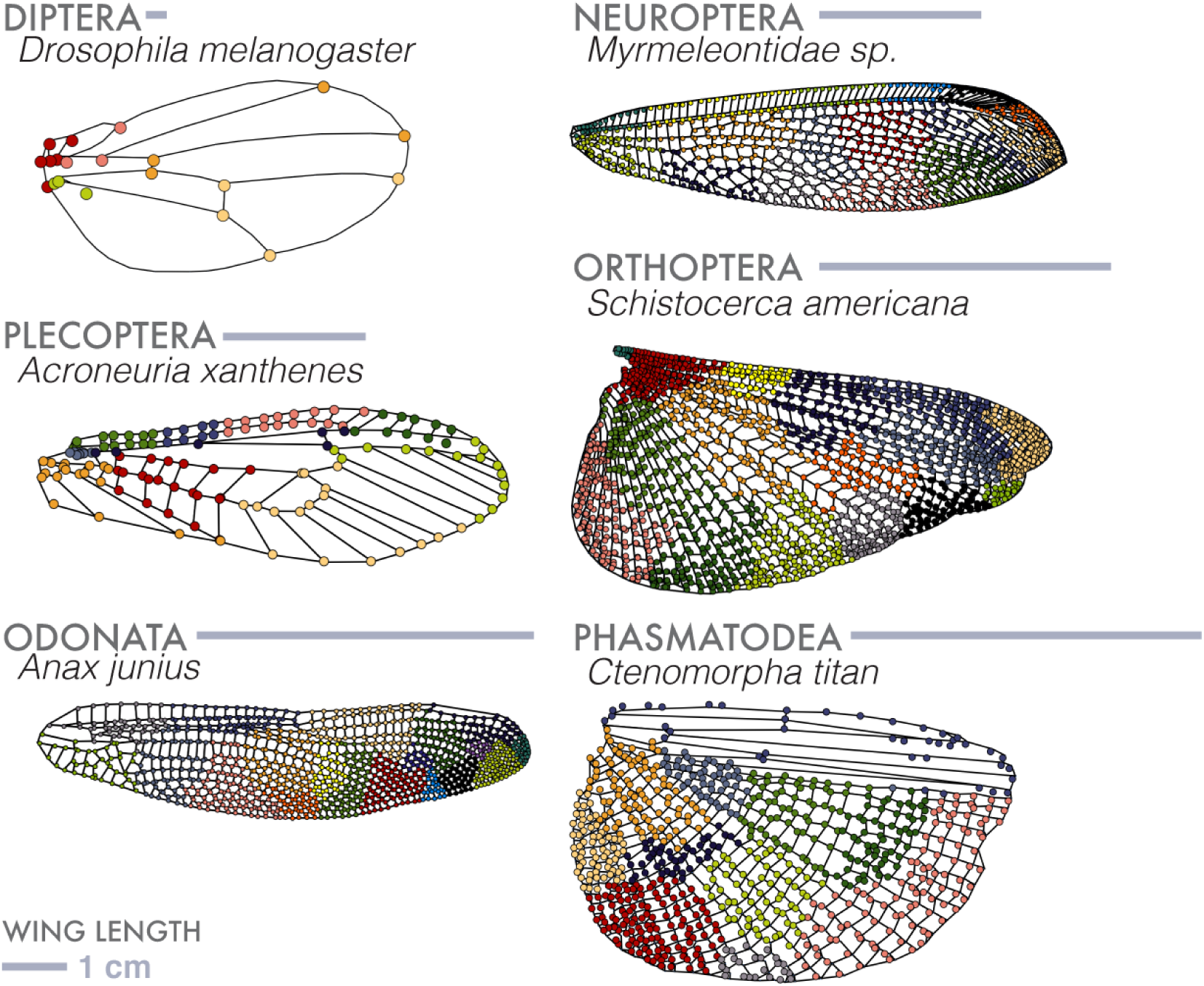
Wing venation network analysis. A wing is a network made up of vein junctions (nodes) and the lengths between them (edges). Using segmented wing images, where each 2D component of the wing is mapped out, we characterize a network using a common community detection algorithm, maximum modularity (see text for details and definitions). Here we show a sampling of wing types and their resultant patterning of clusters.

We deployed simple network analysis to understand venation hierarchies and patterns. deployed. Clustering a network into communities quantifies a third major trait of a wing: a topological measure of the complexity of venation patterns. In Fig. 4 we show range of venation patterns seen in wings, with sparse venation in Diptera at one end, and the dense venation in Odonata at the other end. For wings with sparse venation there are few clusters, e.g. Diptera, Hymenoptera, whereas those with dense venation shows many clusters, e.g. Orthoptera, Odonata. Our comparative network analysis is a first step in understanding how we use venation topology as a precursor to quantifying mechanical and hydrodynamical aspects of wings. It is an open question whether topology metrics will provide insight into wing function.

### Wing domain size and shape distribution

In addition to whole-wing topological and geometric features, we also considered fine-grain features. We found that wings with sparse venation tend to have more rectangular domains (Fig.5a), while wings with dense venation (Fig.5a) tend to have higher numbers of more circular domains. In Figure 5b, we show the domain distributions from representatives of six insect orders with varying complexities of wing shape and venation. This morphospace quantifies domain shapes (circularity, *K*), their distribution in a wing and how much area they occupy within a wing (fractional area, *W*). Within this space, domains at (0, 1) are small and circular while domains at (0,0) are small and rectangular.

**Figure 5.**
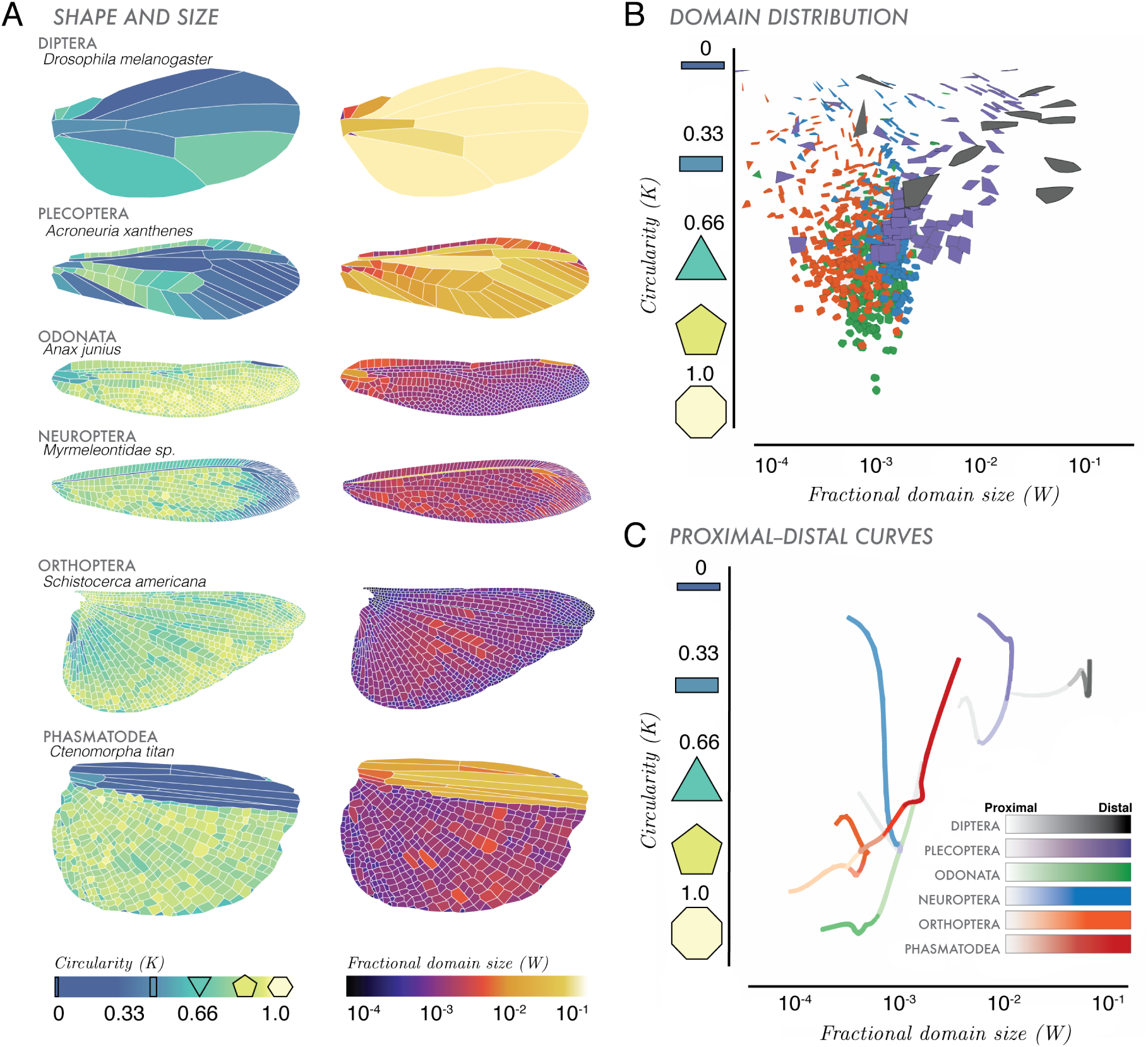
Wing domain sizes and shapes. **(A)** Circularity (*K*) and fractional area (*W*) characterize the distribution of polygonal shapes that make up the vein-bounded domains within wing (see text for definitions). **(B)** Circularity as a function of Fractional Domain Size. **(C)**, Along the proximal to distal (P–D, wing span from base to tip) axis across a wing, we show circularity varies by fading color (lighter = wing base, darker = wing tip), for six insect orders.

Building on the recent work^16^, we plotted domain shapes and how they vary in space across the span of a wing. In Fig. 5c, we consider the proximal to distal axis (P– D axis, wing base to tip) of the wing (similar to wing span). This axis is divided into *N* = 25, rectangular bins, where each bin encompasses all domains across that chord (distance from leading edge and trailing edge of the wing). Following the method of Hoffmann *et al*, we then compute the area-weighted mean area and circularity of all domains within each bin. Then we apply a set of normalized coordinates on this P–D axis, through the computed domain area-circularity space, which is smoothed with a Gaussian of width twice the number of bins (2*/*25 here) and rescaled by the wing’s perimeter. Similarly, portions of the P–D curves located near (0, 1) describe regions of the wing that are dominated by small, circular domains while portions of curves near (1,0) contain domains are characterized by larger, rectangular domains.

In Fig. 5a, we show that domain circularity and fractional area vary within a wing, providing a geometrically minimal description of the internal structure of a wing. Dipteran wings have larger, more rectangular domains, while Odonate wings are made up of smaller, rounder domains. While sparsely veined wings comprise a handful of rectangular domains, e.g. fruit flies (gray, Fig. 5b) and stoneflies (purple, Fig. 5b), other species have thousands of domains. For example, species within the order Megaloptera (Fig. 1) have large, elongate wings, but contain mostly rectangular domains. Furthermore, we see an asymmetry in the distribution of domains, where, as domain number increases, higher numbers of smaller, more circular domains are found in the wing’s trailing edge.

Following the method described in Hoffman *et al.*^16^, in Fig. 5c, we summarize the distribution of domains across the span of a wing, from the proximal to the distal region, showing six species, each of which belongs to a different order. The resultant J-shaped curves represent the entire distribution of domains across the span of a wing. For a dragonfly wing (Odonata, green) the domain morphospace includes more rectangular domains at its wing base (faded green) than at the wing tip, where domains are more circular and more numerous (light green). In contrast, a Plecopteran wing (stonefly, purple), has the opposite domain distribution: larger domains (increased fractional area) and domains are more rectangular at wing tip (dark purple) than at the wing tip. For larger, more elongate wings, rectangular domains tend to be found near the proximal end of the wing while the distal end tends to have smaller, more circular wing domains (Fig. 5a). For approximately 468 Odonate wings (Fig. 5b,c, green), domains near the wing base tend to be rectangular, taking up a larger fractional area of the entire wing. Near the distal end, domains are more circular, taking up less area. In contrast, Neuropteran wings have more elongate and rectangular domains towards the wing tip. Some domains makeup over 10% the total area of the wing (i.e. Diptera), while the smallest account for only 1/10,000 the entire area (i.e. Odonata) of an insect wing. With these domain distributions, our P-D curve morphospace (Fig. 5c) categorizes the spatial geometries of domains across the span of a wing.

## DISCUSSION

Our study has assembled a large dataset of wings from across insect orders. Using this data set, we analyzed segmented wings to create morphospaces comparing simple topological and geometric features of the insect wing, compartmentalizing normalized shape, size and venation structure across the insect phylogeny. By providing these data of segmented images and adjacency matrices, our hope is that others can use different approaches from geometric morphometrics to understand the evolution of wing venation patterns, while also informing modeling efforts for wing flexibility^9,21–23^.

The morphospace of Fig. 3, parses the complexity of wing shape and venation, characterizing contour and internal venation. Results suggest that fliers characterized as forewing dominated tend to have more sparse venation (i.e. Diptera, Hymenoptera) than those that are hindwing or both wing dominated fliers.

The range in shape and branching internal venation also relate to an insect’s flight characteristics and its resistance to damage. Insects with higher numbers of cross veins (thus higher numbers of domains), especially towards the trailing edge, are more likely to reduce tearing and fracturing of the wing that might occur throughout its lifespan^24,25^. Within Odonata, species with larger wings have larger and more numerous domains^16^. While not always applicable across orders, an asymmetry regarding domain number and size across the wing span could be beneficial; from a structural integrity perspective, asymmetry provides fracture toughness^24^. Since cross-veins effectively transfer tensile stresses to neighboring wing domains^25^, wings with higher numbers of small domains (increased cross veins) in the trailing edge could reduce damage propagation^24^.

Our paper is but the first step in addressing the origins and functional consequences of insect wings, and we hope that others will take up the challenges posed by these questions using the datasets and the simple morphometric approaches that we have outlined.

## Supporting information

## Supporting Information (SI)

### SI Datasets

Full methods available in the SI Appendix. This includes: detail on the geometric and topological analysis and lists of species shown in figures. Code: https://github.com/hoffmannjordan/Fast-Marching-Image-Segmentation and https://github.com/hoffmannjordan/size-and-shape-of-insect-wings.

## ACKNOWLEDGEMENTS

The authors would like to thank the librarians at the Ernst Mayr Library, Harvard University for their help in finding insect images, and fruitful discussions with Stacey Combes and Chris **H.** Rycroft.

