## Supplementary material for "Size, shape and structure of insect wings"

**4 Availability of data and code** 5

#### Overview of this document

Here we provide an overview of the data used in the manuscript. We establish the reliability of the collected data, which comes from a wide variety of sources. We then outline the mathematical and computational tools used in the manuscript, all of which are made freely available. Lastly, we provide the algorithms and calculations used in the manuscript.

### 1 Data collection, image processing, and segmentation

The data used in this manuscript comes from a large number of different sources. A few individuals are from original micrographs, however the majority of the data is sourced from previously published sources. While we sampled broadly, our data was limited to those insect wings with low pigmentation. A detailed list of sources is provided in section 2 below.

##### 1.1 Original micrographs

Original micrographs were taken using a CanonScan 9000F Mark II and then processed in `Fiji/ImageJ` using the stitching and extended depth of focus tools.

##### 1.2 Published wing images

Published wing images where either taken (1) from online resources (Biodiversity Heritage Library) or (2) from books at the Ernst Mayr Library, Cambridge, MA. For images taken from online resources, images were contrast-adjusted with Adobe Photoshop when needed. For images taken from textbooks, images were scanned again with the CanonScan.

##### 1.3 Use of images

The supplemental material of [1] show an extensive analysis of comparing insect wing drawings to micrographs. The authors show that these drawings are true-to-form and accurately capture the geometries that we are interested in.

##### 1.4 Segmenting wing images

The segmentation of wing images was done using the code used in [1]. This code is available online at `https://github.com/hoffmannjordan/insect-wing-venation-patterns`. Due to the increased diversity of wing domain shapes, the procedure needed to be modified to handle an increased diversity of wing domains. Specifically, regions that are particularly non-convex need to be properly accounted for. This was done through a combination of code changes (available online) and by manually adding looped regions in certain parts of the wing. These looped regions serve to help in wing domains that are not convex. A loop will not polygonize out, but allows us to add nodes when we polygonize that would not be otherwise captured.

##### 1.5 Vein domain shape and size features for a single wing

After segmenting a wing, each domain is turned into a polygonal representation. This allows us to more accurately compute a variety of statistics about the shape of individual domains, and when grouped, about the shape of the entire wing.

#### 2 Orders and species represented

While our data encompasses 789 wings, we represent several orders with key examples, understanding that within an order the variation of wing venation is highly variable.

**Table 1.** Insects represented in Figure 1

| Species | Family | Order | Reference |
|---|---|---|---|
| *Blattela germanica* | Ectobiidae | Blattodea | [2], Pg. 126, Fig. 199 |
| *Isonychia bicolor* | Isonychiidae | Ephemeroptera | [2], Pg. 265, Fig. 264 |
| *Ogcodes adaptatus* | Acroceridae | Diptera | [3], Fig. 27 |
| *Merope tuber* | Meropiedae | Mecoptera | [2], Pg. 305, Fig. 317 |
| *Anaea archidona (Hewitson)* | Nymphalidae | Lepidoptera | [4], Plate 1, Fig. 1 |
| *Costora iena Mosley* | Sericostomatidae | Trichoptera | [5], Pg. 47, Fig. 23 |
| *Paramigdolus tetropioides* | Vesperidae | Coleoptera | [6], Pg. 140, Fig. 172 |
| *Stylops crawfordi Pierce* | Stylopidae | Strepsiptera | [7], Plate 1, Fig. 5b |
| *Steniolia duplicata* | Crabronidae | Hymenoptera | [8], Plate 1, Fig. 8 |
| *Paracaecilius anareolatus* | Caeciliusidae | Psocodea | [9], Pg. 499, Fig. 1 |
| *Forficula auricularia* | Forficulidae | Dermaptera | [2], Pg. 295, Fig. 305 |
| *Zorotypus hubbardi Caudell* | Zorotypidae | Zoraptera | [10], Pg. 96, Plate 6 |
| *Schistocerca gregaria Forskal* | Locustidae | Orthoptera | [11], Fig. 4 |
| *Eusthenia spectabilis* | Eustheniidae | Plecoptera | [2], Pg. 247, Fig. 246 |
| *Raphidia adnixa* | Raphidiidae | Raphidioptera | [2], Pg. 173, Fig. 168 |
| *Zootermopsis angusticollis* | Termopsidae | Isoptera | [2], Pg. 132, Fig. 126 |
| *Ctenomorpha titan* | Phasmatidae | Phasmatodea | [12], Pg. 122, Fig. 46 |
| *Clothoda nobilis* | Clothodidae | Embioptera | [2], Pg. 265, Fig. 265 |
| *Corydalus primitivus* | Corydalidae | Megaloptera | [2], Pg. 155, Fig. 149 |
| *Cryptoleon nebulosum* | Myrmeleonidae | Neuroptera | [2], Pg. 205, Fig. 202 |
| *Anax junius* | Aeshnidae | Odonata | Salcedo |
| *Mantis religiosa* | Mantidae | Mantodea | [13], Fig. 4 |
| *Leptocysta novatis* | Tingidae | Hemiptera | [14], Pg. 66, Fig. 14 |

#### 2.1 List of species in Figure 1

Table 1 lists species in the Figure 1 phylogeny. Insects hand-caught by MK Salcedo were captured in Bedford, MA at the Concord Field Station in 2015.

#### 2.2 List of species for six representative wings

Table 2 lists the species of our six representative insect wings shown in Figures 2 - 5. The listed Myrmeleontidae sp. was collected by Dino J. Martins in North Kajiado, Oloosirkon, Kenya in 2008.

**Table 2.** Six representative insect wings

| Species | Family | Order | Reference |
|---|---|---|---|
| *Drosophila melanogaster* | Drosophilidae | Diptera | [15], Pg. 166 |
| *Acroneuria xanthenes Newm.* | Acroneurinae | Plecoptera | [16], Plate 16, Fig. 10 |
| *Anax junius* | Anax junius | Odonata | Salcedo |
| *Myrmeleontidae sp* | Myrmeleontidae | Neuroptera | Martins |
| *Schistocerca americana* | Acrididae | Orthoptera | Salcedo |
| *Ctenomorpha titan* | Phasmatidae | Phasmatodea | [12], Pg. 122, Fig. 46 |

### 3 Quantitative characterization

#### 3.1 Initialization

For each polygonized wing, the coordinates are rescaled such that the entire wing has area 1. This effectively removes all size information for each wing, allowing comparison between species. In each of the analyses discussed below, the area is first rescaled before applying the technique [1].

#### 3.2 Network analysis

We deployed a suite of network analysis tools on the diverse set of insect wing geometries. We tried both weighted [17] and unweighted analysis [18], though overall ground the results difficult to interpret. Future work, with more quantitative mechanical measurements or knowledge of fluid movement could easily extend this work.

##### 3.2.1 Unweighted

After polygonizing an insect wing, we construct a list of vertices. We then construct an adjacency matrix, $M = N \times N$ matrix of 0's, where $N$ represents the number of vertices. For each $i, j$ pair of vertices that are connected, we set $M_{i,j} = 1$.

##### 3.2.2 Weighted

Applying weighted connections between nodes (vein junctions) with length, $L$, allowed weighting $M_{i,j} = 1/L^n$ where $L$ is the distance between nodes $i$ and $j$ and $n = 1, 2$. We also looked at an analysis using the resistance between nodes, $L/r^4$, where $r$ was computed using the 2D width from segmented images of original micrographs (giving us an approximate thickness of veins). Rather than measure the radius of each vein segment, we took a handful of original micrographs and measured approximately 50 cross vein radii and their location. We then interpolated over the wing as a proxy.

#### 3.3 Curvature

When computing curvature [19], $\kappa$, we do not use our polygonized wing. Instead, we use our original segmentation and extract the boundary of the wing region. Then we orient each wing such that the base of the wing is on the left and the perimeter of the wing runs clockwise. We choose a distance of $N = 0.02L_P$, and for each point on the perimeter, $P$, we take $p_i$, $p_{i-N}$ and $p_{i+N}$ where $L$ is length of the entire perimeter. From here, we perform a linear-least square fit to a circle, where we define $\kappa = 1/R$.

#### 3.4 Internal venation length

For all nodes $N$, we sum up the distances between all nodes. In doing this calculation, we then subtract off the perimeter, which we calculate separately.

##### 3.4.1 Proximal-to-Distal morphology traces

We characterize representatives of our large dataset using the P–D morphology traces introduced in [1]. We divide the wing into 25 equal spaced bins along the long axis of the insect wing. To calculate the mean circularity of each slice $i$, which we denote $\mathcal{S}_i$, we compute

$$\frac{1}{\sum_{\mathcal{P}_j} \text{Area}(\mathcal{S}_i \cap \mathcal{P}_j)} \sum_{\mathcal{P}_j} \text{Area}\,(\mathcal{S}_i \cap \mathcal{P}_j)\, \mathcal{C}(\mathcal{P}_j) \tag{1}$$

where the sum is applied over all polygons $\mathcal{P}_j$. This produces a smoothly varying mean circularity as we move along the long axis of the wing from base to wing tip. For polygon $\mathcal{P}_i$, we obtain the area fraction $f_i$ that overlaps with the bin. We construct the vector of all area fractions in bin $i$, denoted $\vec{F}_i$. We also have the vector of all circularities $\vec{C}$ and all areas $\vec{A}$. For each bin, we compute

$$(\vec{F}_i \cdot \vec{A}, \vec{F}_i \cdot \vec{C}) \tag{2}$$

giving us the weighted mean area and the mean circularity of the wing domains in the region.

### 4 Availability of data and code

Integer `csv` files of segmented images of all data along with code used in the manuscript are freely available. All code and data can be found at https://github.com/hoffmannjordan/size-and-shape-of-insect-wings.
